# One size does not fit all: Caste and sex differences in the response of bumblebees (*Bombus impatiens*) to chronic oral neonicotinoid exposure

**DOI:** 10.1101/351809

**Authors:** Melissa W. Mobley, Robert J. Gegear

**Affiliations:** Department of Biology and Biotechnology, Worcester Polytechnic Institute, Worcester, 01609-2280, U.S.A

## Abstract

Neonicotinoid insecticides have been implicated in the rapid global decline of bumblebees over recent years, particularly in agricultural and urban areas. While there is much known about neonicotinoid toxicity effects at the colony stage of the bumblebee annual cycle, far less is known about such effects at other stages critical for the maintenance of wild populations. In the present work, individual-based feeding assays were used to show that chronic consumption of the widely used neonicotinoid clothianidin at a field-realistic average rate of 3.6 and 4.0 ng/g·bee/day reduces survival of queen and male bumblebees, respectively, within a 7-day period whereas consumption at a similar rate of 3.9 ng/g·bee/day in workers had no effect on survival. To test the hypothesis that males have a lower tolerance for oral clothianidin exposure than workers due to their haploid genetic status, RNAseq analysis was used to compare the transcriptomic responses of workers and males to chronic intake of clothianidin at a sub-lethal dose of 0.37ng/day for 5 days. Surprisingly, only 25/100 clothianidin-induced putative detoxification genes had expression levels that differed in a sex-specific manner, with 17 genes showing increased expression in workers. Sub-lethal clothianidin intake also induced changes in genes associated with a variety of other major biological functions, including locomotion, reproduction, and immunity. Collectively, these results suggest that chronic oral toxicity effects of neonicotinoids are greatest during mating and nest establishment phases of the bumblebee life cycle. Chronic oral toxicity tests on males and queens are therefore required in order to fully assess the impact of neonicotinoids on wild bumblebee populations.

## Introduction

Bumble bees have rapidly declined in abundance, species richness, and geographic distribution on a global scale over recent years [1, 2]. In North America alone, nearly half of the bumblebee species have reached historically low numbers [3-5], including one species, *Bombus affinis*, which was recently listed as an endangered species by the U.S. Fish and Wildlife Service. From an ecological perspective, these declines pose a significant threat to the function and diversity of temperate ecosystems due to the critical keystone role that bumblebees play as pollinators of native flowering plants. Reports of parallel reductions in bee-pollinated plant species suggest that pollinator decline-mediated effects on wildlife diversity at higher trophic levels may already be well underway [1, 6, 7]. It is therefore imperative that all anthropogenic stressors contributing to species decline in bumblebees be identified and mitigated as soon as possible.

One stressor thought to contribute to wild bumblebee decline in urban and agricultural areas is a newly developed class of pesticides called ‘neonicotinoids’ [8]. From a pest management perspective, neonicotinoids are a highly effective form of insect control because they specifically target nicotinic acetylcholine receptors in the insect central nervous system, leading to paralysis and death [9]. Unlike most other pesticide classes, neonicotinoids are also systemic, meaning that they are readily taken up by the roots and distributed throughout the entire plant, thereby protecting it from pest damage over the entire growing season [10].

Despite these benefits, neonicotinoids pose a significant threat to beneficial insects such as pollinators because they are also translocated to floral nectar and pollen, presenting a route of oral exposure. Of particular concern to wild pollinator populations is the fact that neonicotinoids can be transported away from the area of application to adjacent natural areas [11, 12] and then contaminate wildflower resources [2, 13, 14]. For example, clothianidin, one of the newest and most potent neonicotinoid formulations, has been detected in numerous wildflower species heavily visited by insect pollinators at concentrations ranging from 4 to 215 ppb [2, 14]. Compounding this issue, neonicotinoid residues can persist in the soil for years after a single application [15-17], and increase in concentration with repeated annual application [18]. Wild bumblebees and other insect pollinators therefore have the potential to be orally exposed to neonicotinoids at many life stages occurring outside of the blooming period and geographic location of the targeted plant species.

Yet, the overwhelming majority of neonicotinoid toxicity studies on bumblebees to date have focused only on the colony stage, relying on metrics such as worker survival, larval growth, and reproductive output to estimate the potential impact of exposure on wild populations [19]. While such colony-focused risk assessment protocols may be adequate in the context of crop pollination, determining the impact of widespread neonicotinoid use on threatened bumblebee species in an ecological context requires more comprehensive assessment protocols that consider potential impacts on queens and males, whose survival during mating, overwintering, and nest establishment stages of the life cycle has a direct effect on population stability (Fig 1).

**Fig 1.**
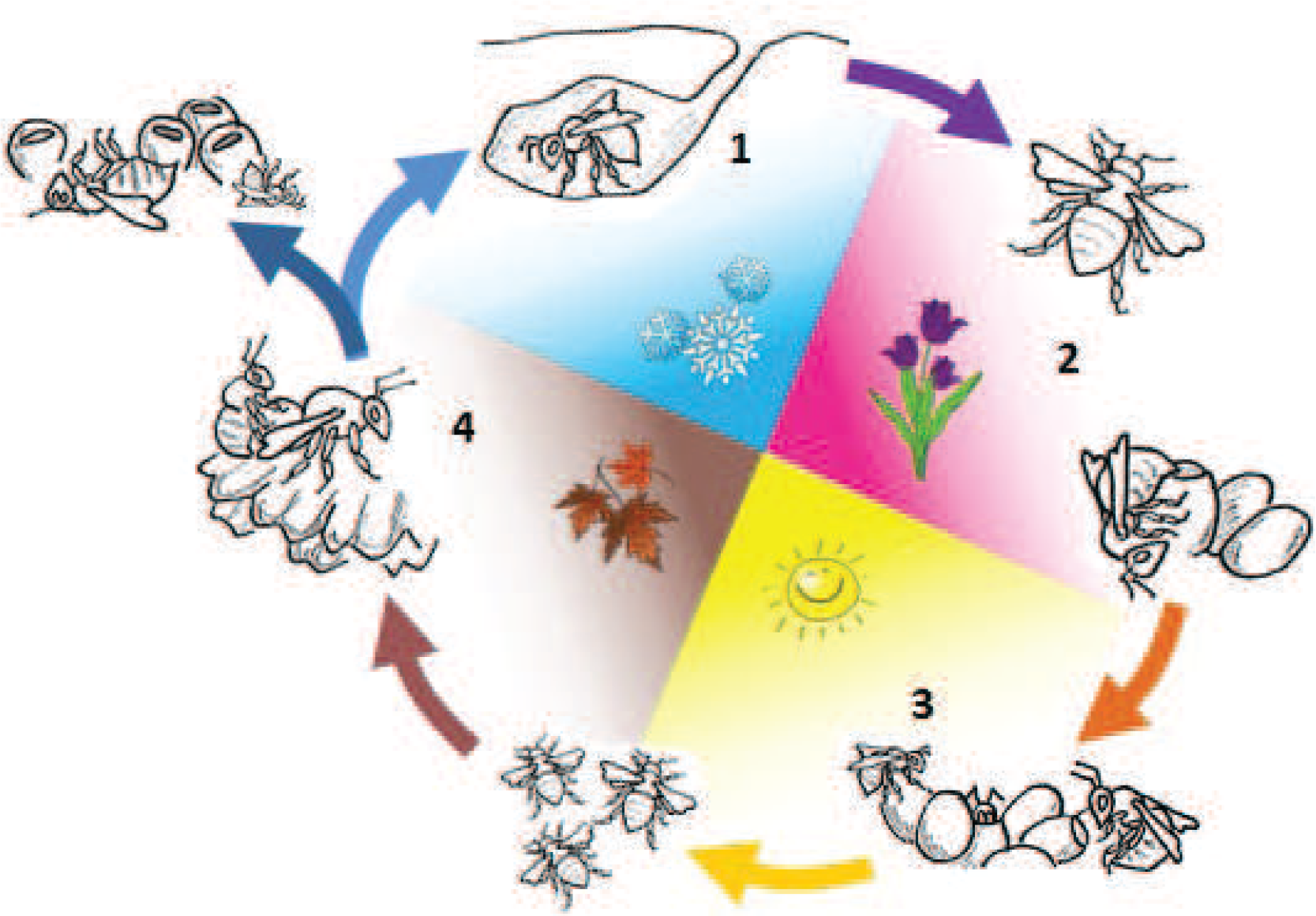
Annual cycle of social bumblebees. (1) Mated queens emerge from their overwintering site in the spring and forage while search for a suitable nest site. (2) Queens collect floral resources and store them in the nest until the first brood of worker bees emerges. (3) Workers forage for the colony so that it can continue to produce more workers and later, queens and males (reproductives). (4) Virgin queens and males leave the nest to feed and search for mates. Once mated, queens, which are the only individuals to survive through the winter months, continue to feed until they find a suitable overwintering site where they reside until the spring, thus completing the cycle.

Several morphological and physiological characteristics of queen, worker, and male bumblebees suggest that individuals from each group may vary in their capacity to cope with chronic oral neonicotinoid exposure. For example, males and workers have a smaller body size than queens and therefore may experience mortality effects at lower neonicotinoid concentrations, as has been shown at the species level [20]. Males are also haploid whereas queens and workers are diploid, which may limit gene products available for males to detoxify neonicotinoids. Such ‘haploid susceptibility’ is known to occur in the context of immunocompetence [21], but has yet to be studied in the context of metabolic resistance to pesticides. Alternatively, males and queens have greater energetic demands due to their reproductive status [22, 23] and therefore may have fewer resources available for detoxification processes than sterile workers.

To explore these possibilities, the present study uses a novel individual-based feeding assay to directly compare the mortality response of queen, worker, and male *Bombus impatiens* to chronic consumption of the widely used neonicotinoid clothianidin at field-realistic concentrations of 5-10ppb. Assaying bumblebee test populations at the individual level provides a much more robust estimate of chronic lethality level than more traditional assays on small groups (e.g. microcolonies) because drug intake rates can be directly monitored in all test individuals and then easily standardized by adjusting for individual variation in body size [24]. Our feeding experiments revealed that chronic lethality effects of clothianidin are much greater in males than workers even when controlling for differences in body size. To test the potential role of male haploidy in driving this sex difference, we then used RNAseq analysis to compare the transcriptomic responses of workers and males to consumption of clothianidin at a sub-lethal daily dose over a 5-day period. Following previous work on bumblebees by [25], putative genes related to clothianidin detoxification were identified and classified based on the two well-characterized xenobiotic detoxification phases in insects, with Phase 1 processes metabolizing the xenobiotic to less toxic compounds through oxidation, hydrolysis, and reduction reactions, and Phase 2 processes breaking down the metabolites produced from Phase 1 processes through conjugation reactions. Based on the findings of [25] that another bumblebee species (*B. huntii*) has fewer constitutively expressed detoxification genes than workers, it was predicted that male *Bombus impatiens* would have fewer and/or lower expression of inducible detoxification genes compared to workers.

## Results

### Chronic oral toxicity effects of clothianidin differ among queen, worker, and male bumblebees

Chronic consumption of clothianidin at field-realistic concentrations increased mortality rates in queen, worker, and male test populations (Fig 2). However, clothianidin concentrations needed to produce such effects varied in a caste- and sex-dependent manner. Table 1 shows intake rates reducing survival in queens, workers, and males controlling for differences in body size and daily clothianidin dose. Compared to 0ppb controls, queens consuming clothianidin at a concentration of 10ppb (mean intake rate ± SD = 3.61±0.71 ng/g·bee/day) had reduced survival over the 7-day test period (X^2^=15.6, df=1, P<0.0001). While workers orally exposed to clothianidin at a concentration of 10ppb (mean intake rate ± SD = 5.48±1.95 ng/g·bee/day) also had reduced survival (X^2^=54.7, df=1, P<0.0001), with the test population reaching 50% mortality at day 5. Interestingly, worker consumption mean (±SD) intakes rates of 3.85 (±1.6) ng/g·bee/day (7ppb solution) had no effect of survival, indicating that workers are better able to cope with chronic oral clothianidin exposure than queens when controlling for consumption rate and body size.

**Fig 2.**
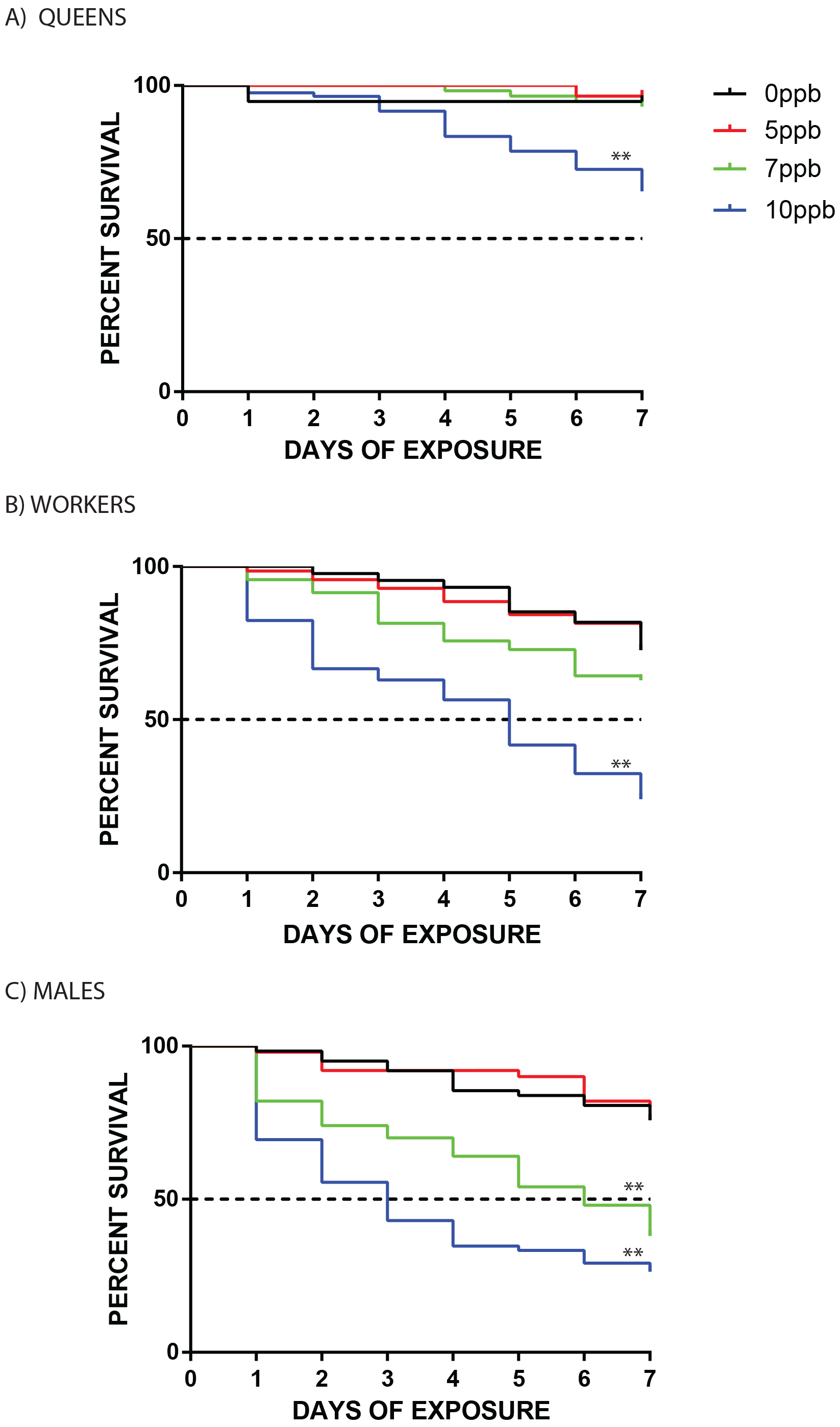
Kaplan-Meier survival plots for *Bombus impatiens* (A) queens, (B) workers, and (C) males, orally exposed to test solutions containing different concentrations of clothianidin daily for 7 days. Black = 0ppb; red= 5 ppb; green = 7 ppb; blue = 10ppb. A Mantel-Cox Log-rank analysis was used to test for differences in survival between each treatment group relative to the 0ppb control group over the 7-day period. **, p<0.0001.

**Table 1.** Response of queen, worker, and male bumblebees (*Bombus impatiens*) to chronic clothianidin exposure controlling for differences in body size and daily dose. Workers and males consumed 75 μL of solution per day and due to their larger size, queens consumed 200 μL of each solution per day. Yellow shading = no effect on survival; blue shading = reduced survival. All values are shown as mean ± SD.

|  | Concentration of clothianidin solution |  |  |  |  |  |
| --- | --- | --- | --- | --- | --- | --- |
|  | 5ppb(ng/mL) |  | 7ppb(ng/mL) |  | 10ppb(ng/mL) |  |
|  | Body mass (g) | Intake rate (ng/g.bee/day) | Body mass(g) | Intake rate (ng/g.bee/day) | Body mass (g) | Intake rate (ng/g.bee/day) |
| **Queens** | 0.58±0.1<br>3 | 1.79±0.39 | 0.59±0.1<br>1 | 2.45±0.48 | 0.57±0.1<br>1 | 3.61±0.71 |
| **Workers** | 0.16±0.0<br>5 | 2.60±0.83 | 0.15±0.0<br>4 | 3.85±1.6 | 0.15±0.0<br>4 | 5.48±1.95 |
| **Males** | 0.14±0.0<br>4 | 2.94±0.86 | 0.14±0.0<br>4 | 4.01±1.08 | 0.13±0.0<br>3 | 6.05±1.35 |

Similar to workers and queens, males consuming the 10 ppb clothianidin solution (mean intake rate ± SD = 6.05 ±1.35 ng/g·bee/day) had reduced survival compared to the 0ppb control (X^2^=38.8, df=1, P<0.0001); however, the test population reached 50% mortality after just 3 days. In addition, males consuming clothianidin at a concentration of 7 ppb (mean intake rate ± SD = 4.01 ±1.08 ng/g·bee/day) showed reduced survival compared to 0ppb controls (X^2^=17.7, df=1, P<0.0001), with the test population reaching 50% mortality at day 6. The vast majority of queens, workers, and males fully consumed the clothianidin solutions during testing (Table S1), indicating that differences in mortality were caused by differences in the toxicity effects of clothianidin rather than starvation (anti-feeding behavior).

### Sex differences in the transcriptomic response of bumblebees to sub-lethal doses of clothianidin

RNAseq analysis revealed that chronic oral exposure to clothianidin at a sub-lethal concentration of 5ppb for 5 days affected expression of 147 genes in workers and 200 genes in males. Of the 347 pooled clothianidin-induced genes (147 worker, 200 male), only 15 were present in both sexes, yielding a total of 332 unique clothianidin-induced genes. Of these genes, 100 were classified as having a putative role in detoxification and the other 232 were associated with other biological functions (Tables 2, S2, and S3).

**Table 2.**
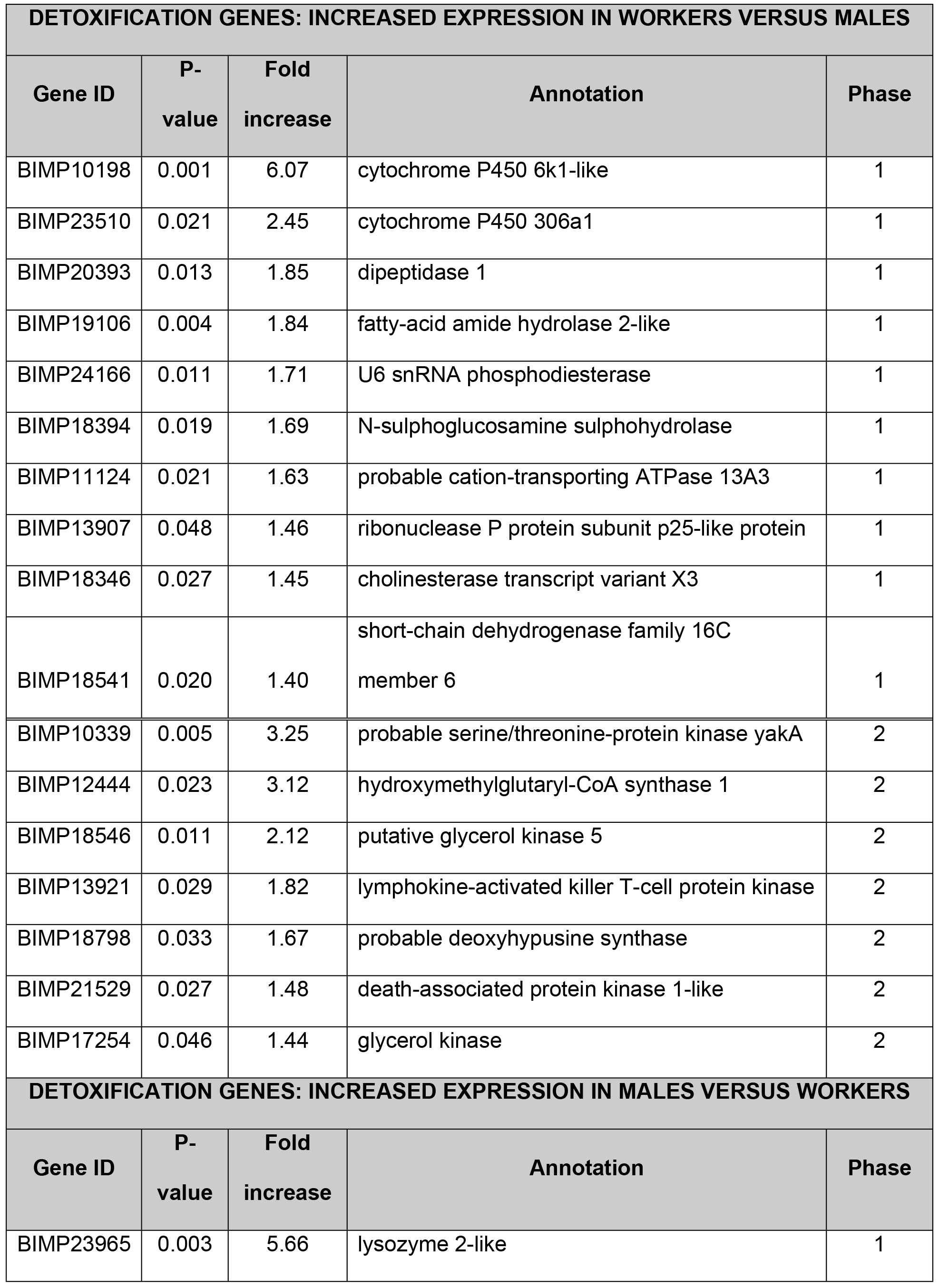
Sex-specific expression of inducible detoxification genes in bumblebee workers and males orally exposed to clothianidin at a sub-lethal daily dose over 5 consecutive days. Phase 1 detoxification processes include oxidation, hydrolysis, and reduction reactions while Phase 2 processes include conjugation reactions. Expression levels were determined through RNAseq analysis. Upper portion = genes with increased expression in workers; lower portion = genes with increased expression in males. Clothianidin-inducible genes were initially identified by comparing transcriptomic expression patterns between individuals fed either 5ppb clothianidin solution or the same volume of solution containing 1.4% DMSO (vehicle control) solution separately for each sex and then pooling genes with significantly altered expression together.

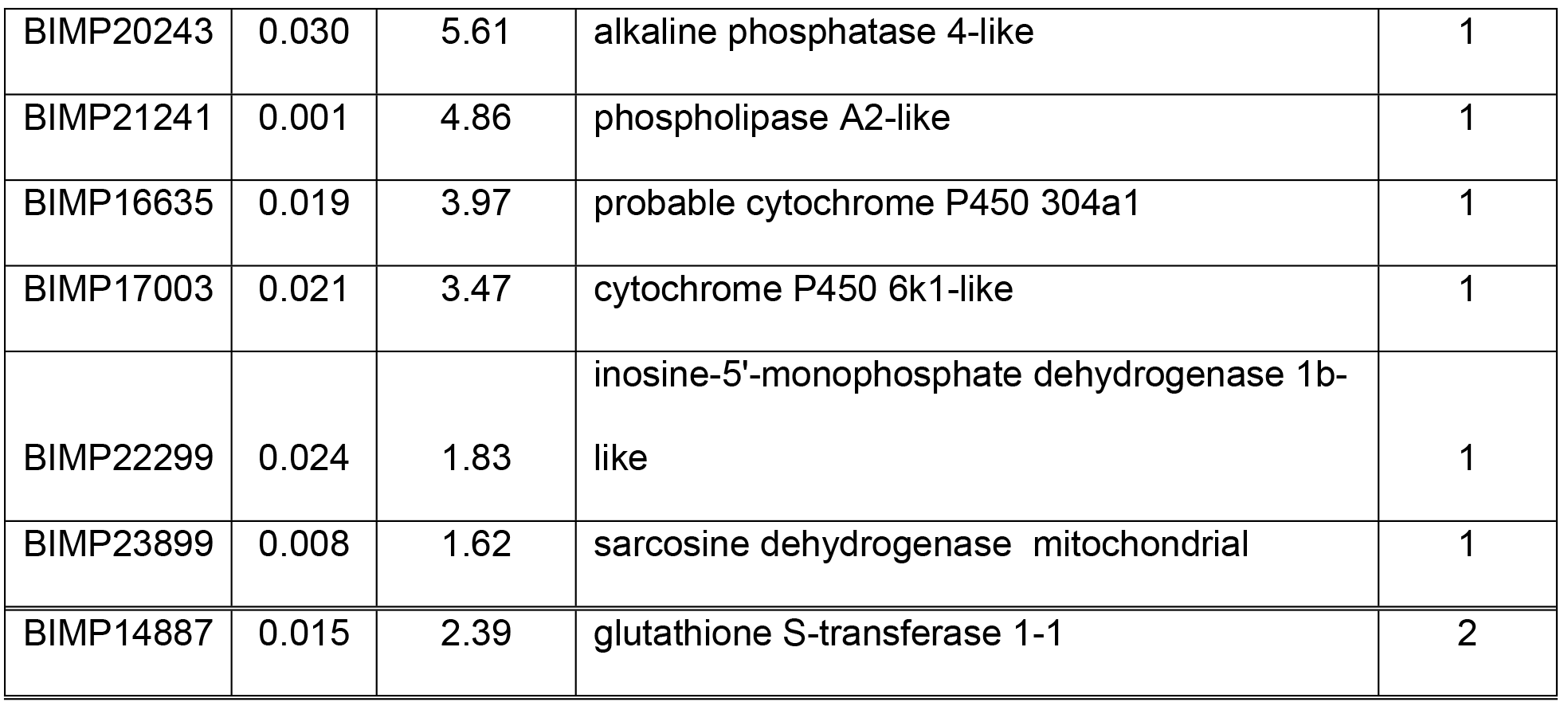

Only 77 of the 332 clothianidin-sensitive genes were differentially expressed between workers and males: 48 genes had higher expression in workers and 29 genes had higher expression in males. Of these 77 genes, only 25 were associated with detoxification processes 17 (10 Phase 1 and 7 Phase 2) had higher expression in workers and 8 (7 Phase 1 and 1 Phase 2) had higher expression in males (Table 2; Fig 3). Of the remaining 52 genes associated with biological functions other than detoxification, 31 genes had higher expression in workers and 21 genes had higher expression in males. Biological functions, fold changes, and accession numbers for these 52 genes are shown in Table S2. Although many of these genes may play some as yet unknown functional role in detoxification, currently known biological functions include immunity, learning and memory, reproduction, and signal transduction. We also found eight genes whose function we could not identify based on current databases. Information for the 255/332 clothianidin-inducible genes with conserved expression between workers and males are provided in Table S3.

**Fig 3.**
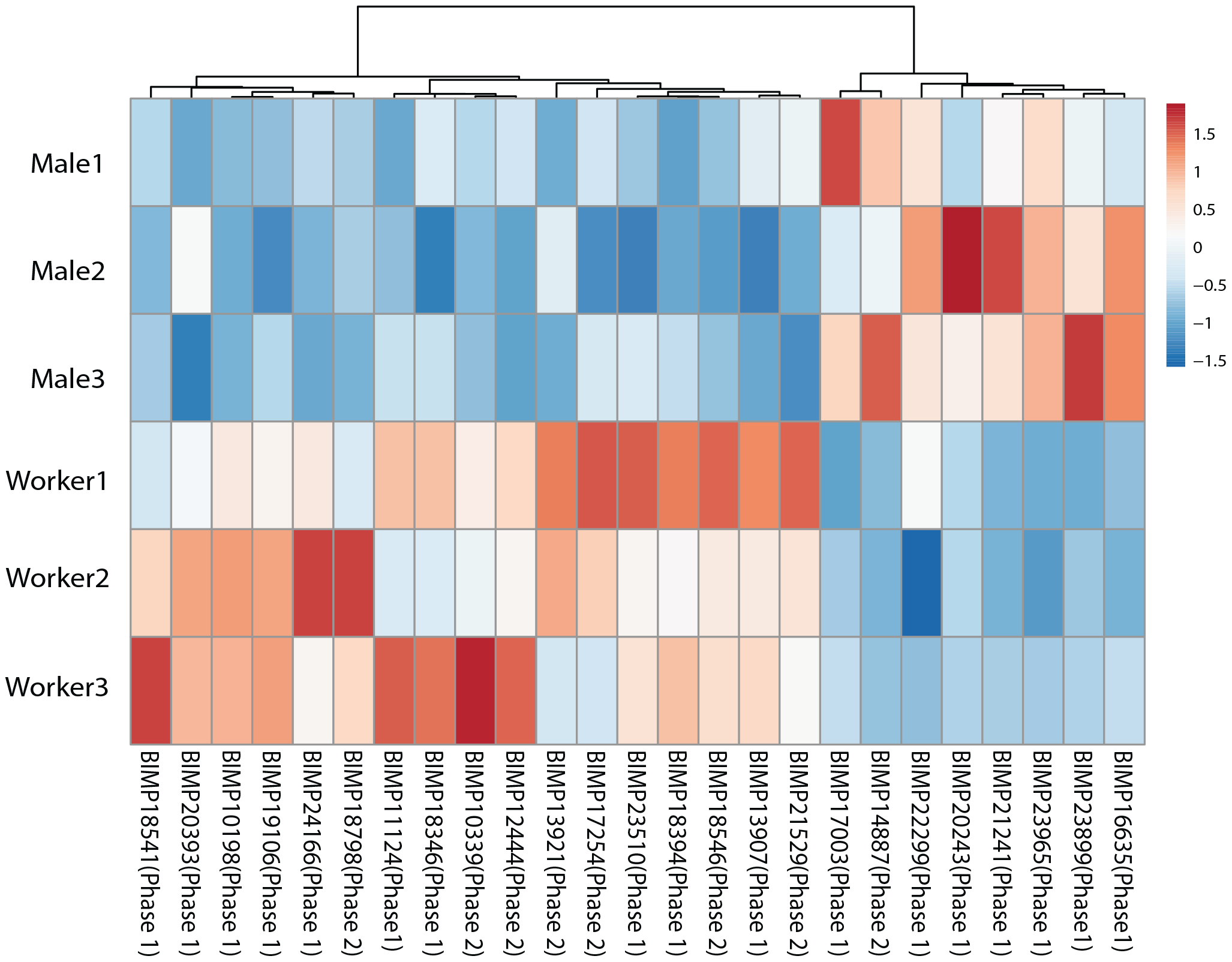
Heatmap showing the 25 putative detoxification genes that were differentially expressed between workers and males after 5 consecutive days of oral exposure to clothianidin. Phase 1 detoxification processes include oxidation, hydrolysis, and reduction reactions while Phase 2 processes include conjugation reactions. Legend on the right shows relative fold change corresponding to each color.

## Discussion

The present study shows that chronic consumption of field-realistic doses of the widely used neonicotinoid pesticide clothianidin has differential effects on the survival of queen, worker, and male bumblebees. Controlling for body size, queens and males consuming clothianidin at an average rate of 3.6 and 4.0 ng clothianidin/g·bee/day, respectively, had reduced survival over a 7-day period whereas a similar average consumption rate in workers (3.9 ng/g·bee/day) had no mortality effect. Although clothianidin reduced queen and male survival at similar daily consumption rate, only male test populations reached 50% mortality over the testing period, indicating that chronic oral lethality effects for clothianidin are slightly greater in males than queens. The observed mortality response of all test groups to chronic clothianidin consumption cannot be attributed to a starvation effect as food avoidance and vomiting behaviors were not observed during the testing period. Collectively, these findings demonstrate that queens and males are much more vulnerable to clothianidin toxicity effects than workers, highlighting the importance of incorporating queen and male survival assays into current colony-focused ‘higher tier’ protocols for assessing the potential risk of various neonicotinoid formulations to wild pollinator populations [19]. In addition, the observed reductions in survival in *B. impatiens* workers chronically consuming clothianidin at a concentration of 10ppb is notably less than the 20ppb chronic lethality threshold reported for worker honeybees [26] as well as the 25ppb threshold reported for workers of the European bumblebee species *B. terrestris* [27], indicating that chronic oral lethality effects of clothianidin are relatively high in *B. impatiens*. Our findings thus add to the rapidly growing body of evidence that neonicotinoid oral toxicity effects can vary considerably among insect pollinator species [28-31].

In addition to increasing mortality, chronic oral exposure to clothianidin over a short time period (5 days) was found to have profound sub-lethal effects on workers and males at the genomic level, altering expression of a total of 332 genes associated with a wide variety of biological functions, including immunity, neuronal signal transduction, locomotion, reproduction, and several fundamental cellular processes (Tables 2, S2, and S3). These results are consistent with a growing number of studies at the organismal level showing that neonicotinoids can have substantial sub-lethal effects on pollinators [32-36]. For example, clothianidin consumption induced changes to genes associated with reproductive output in males such as outer dense fiber protein 3-like and lutropin-choriogonadotropic hormone receptor-like (Table 2), which is consistent with previous work showing that oral exposure to imidacloprid reduced sperm viability in males bees by 50% [37]. Under natural conditions, chronic exposure to low doses of clothianidin in wildflower nectar thus has the potential to influence the dynamics of wild bumblebee populations by reducing male and queen reproductive capacity and at slightly higher doses, reducing the number of mating individuals in the fall and the number of queens establishing nests in the spring.

Somewhat surprisingly, the genomic response of workers and males to sub-lethal clothianidin doses did not provide unequivocal evidence that the haploid genetic status of males renders them less able to cope with oral clothianidin exposure. Of the 100 putative detoxification genes that were affected by clothianidin, 75 had a similar expression level in workers and males. These similarly expressed genes included many ‘classic’ detoxification genes such as cytochrome P450s, glutathione S-transferases, and carboxylesterases [38]. There was, however, a strong sex bias in direction predicted by the haploid susceptibility hypothesis for the 25 remaining clothianidin-induced detoxification genes that differed between the sexes. Compared to males, workers had higher expression in more than twice genes associated with Phase 1 hydrolysis reactions and remarkably, higher expression in seven times as many genes associated with Phase 2 conjugation reactions. The fact that workers and males had a similar number of highly expressed Phase 1 genes related to oxidation-reduction reactions suggests that are were equally capable of initially breaking down clothianidin into intermediate metabolites, but differed in the capacity to subsequently transform such metabolites into non-toxic compounds (Phase 2 reactions). These results are similar to those reported for constitutively expressed detoxification genes in *B. huntii* [25], which is closely related to *B. impatiens*. However, it must be noted that Xu et al. identified over 250 constitutively expressed genes for conjugation compared to the 18 inducible genes found in the present study. Thus, despite the fact that there were substantial differences in the mortality response to oral clothianidin exposure, they were not strongly correlated with the number and expression of inducible detoxification genes. Of course it is possible that haploidy in males results in the reduced expression of few key genes in the clothianidin detoxification pathway, but further genetic manipulation experiments are required to determine whether or not this is the case.

The present work demonstrates the potential conservation benefits of using a genomic approach to quantify exposure of individual pollinators to extremely low doses of neonicotinoids, an idea that has been recently advocated by others [39]. Currently, one of the major problems with studying neonicotinoid effects on wild bee populations is that individual tissue samples have concentrations that are too low to be quantified with a high level of accuracy and reliability using conventional methodology and instrumentation. Indeed, the main laboratory at the United States Department of Agriculture responsible for measuring neonicotinoid levels in tissue samples was unable to detect clothianidin in our 5ppb in test solution, yet it produced a robust genetic response in both males and workers. To address this problem, the current study has identified a set of clothianidin-induced genes that could potentially be used to develop highly sensitive ‘eco-tools’ for assessing neonicotinoid exposure wild bumblebee populations. Such tools could also be used to gauge sub-lethal effects on behavior (e.g. reduced cognitive function) and physiology (e.g. reduced sperm production), which are often difficult to assess in the field at the organismal level. Comparative genomic approaches would also provide unique insight into why chronic oral toxicity levels for neonicotinoids vary considerably among (and within) wild bees species [31], greatly accelerating efforts to determine the impact of neonicotinoid use on wild pollinators and the global biodiversity that they ultimately support.

## Materials and Methods

### Experimental Design

#### Bumble bees

Virgin queens, workers, males were obtained from commercial *Bombus impatiens* colonies (Biobest Biological Systems, Leamington, Canada). Colonies initially contained approximately 100 workers and were subsequently supplied with 30% (stock) sugar solution (Grade A pure honey mixed with distilled water to the desired concentration, measured with a Bellingham & Stanley hand-held refractometer (Suwanee, USA)) and wildflower pollen *ad libitum* to facilitate continued production of workers and later production of queens and males. Multiple colonies were used across experimental trials to control for potential inter-colony differences in clothianidin sensitivity.

For chronic oral toxicity tests, queen, worker and male bees were collected and placed in a 4°C refrigerator. Once immobile, individuals were weighed to the nearest milligram, marked with acrylic paint for identification, and then placed in a 16oz plastic housing container with a screen lid for ventilation (Fig S1). Each lid had a circular opening with a removable plastic plug that was positioned directly above a plastic ‘feeding bowl’ fastened to the bottom of the container.

In this way, test solutions could be dispensed into the bowls with minimal disturbance to the bee. Each container was also supplied with 1-1.5 grams of a pollen paste (a dough-like mixture of and stock sugar solution), which was replaced as necessary.

All housing containers were kept in Percival Scientific (Perry, USA) environmental chambers set to 22-25°C, 50 % humidity, and a 12 hour light/dark cycle. Bees were fed 75 μL (workers and males) or 200 μL (queens) of untreated stock sugar solution once daily within 3 hours of the start of the light cycle. Preliminary experiments with stock sugar solution determined that these volumes are completely consumed by individuals over a 24-hour period while having no effect on survival or activity level. All handling of housing containers was conducted under red light in order to minimize additional stress on test bees.

#### Clothianidin Solutions

Analytical-grade clothianidin (Sigma-Aldrich, USA,) was dissolved in dimethyl sulfoxide (DMSO), as has been done in other neonicotinoid toxicity studies [40], to a concentration of 5 × 10^6^ ng/mL or ppb (parts per billion) and serially diluted down to 5 × 10^2^ or 5 × 10^1^ ppb. Diluted clothianidin solution was added to stock sugar solution to create feeding solutions of 10, 7, and 5 ppb (ng/mL). Concentrations of clothianidin selected were well within the range present under field conditions [41]. Drug mixtures were prepared within 2 days of the start of a trial, stored in conical tubes in the dark at 4°C and only taken out for immediate use.

To confirm that test solutions contained the correct amount of clothianidin, fresh samples of 0, 5, 7, and 10 ppb test solutions (were sent to the U.S.D.A. Agricultural Marketing Service’s National Science Laboratories Testing Division (Gastonia, USA). Clothianidin residues were measured using Mass Spectrometry with a LOQ of 5 +/- 1 ppb. Although the 5ppb test solution was below the minimal level of detection by the U.S.D.A. equipment, 7 and 10 ppb test solutions (3 technical replicates each) yielded mean values (+/-SE) of 4.1 +/- 3.5 and 7.3 +/-1.4 ppb, respectively.

### Test Procedure for feeding experiments

Prior to each trial, all bees were fed stock sugar solution at the previously described volumes daily until there were no observed deaths over a 48-hour period. Individuals not consuming all of the sugar solution within a 24 hour period were also removed. This process ensured that bees were in good health prior to testing.

After the acclimation period, individuals were assigned to either the 0 ppb (control), 5 ppb, 7ppb, or 10ppb treatment group, with colony origin and bee size balanced among groups. Location of housing containers within and between shelves inside the environmental chamber was randomized in order to control for potential positional effects on survival. Total number of bees tested at each clothianidin concentration was as follows: 208 at 0ppb (88 workers, 58 queens, 62 males), 178 at 5ppb (70 workers, 58 queens, 50 males), 178 at 7 ppb (70 workers, 58 queens, 50 males), and 264 at 10 ppb (108 workers, 84 queens, 72 males). These numbers were obtained through 11 independent worker, 8 queen, and 12 male trials, with bees in each trial obtained from at least two colonies. Given queens required a greater volume of test solution to remain active than workers and males due to their larger body size, we expressed exposure level for each group as ng clothianidin consumed per gram of bee per day (ng/g·bee/day). Mean (±SD) mass in grams for queens = 0.58±0.12 g; workers = 0.15±0.05 g; males = 0.14±0.04 g. In this way, sex and caste differences in clothianidin sensitivity could be established while controlling for differences in body size and consumption rate.

Each day over a 7-day testing period, individual housing containers were removed from the environmental chamber within three hours after to the beginning of the light cycle, and the number of dead bees was recorded. In the rare case that a test individual was not dead but showed inverted orientation, extremity spasms, and unresponsiveness, it was recorded as dead and euthanized. Presence/absence of test solutions was also checked daily and feeding bowls were replenished. Immediately following data collection, bees were returned to the environmental chamber.

To confirm that consumption of dimethyl sulfoxide (DMSO) at the concentration used in experiments had no effect on survival of queens, workers and males, additional feeding trials were conducted with stock sugar solution containing 1.4% DMSO. As in the clothianidin test trials, workers and males were fed 75 μL of solution per day and queens were fed 200 μL/day over a 7-day period. Survival in the control (0ppb) group (see above for numbers in each group) was then compared to survival in the 1.4% DMSO group for queens (24 treated individuals), workers (55 treated individuals), and males (87 treated individuals). Results of Mantel-Cox Log-rank analyses revealed that consuming 1.4% DMSO for seven days had no effect on survival of queens (X^2^=0.61, df=1, P=0.43), workers (X^2^=0.26, df=1, P= 0.60) and males (X^2^=0.08, df=1, P=0.77).

#### Statistical Analysis

Survival data from each caste was compiled over the full 7-day clothianidin trial period. All remaining survivors were censored. For each clothianidin concentration, Kaplan-Meier survival curves [42] were generated for queens, workers, and males in GraphPad Prism 6 (La Jolla, USA) and a Mantel-Cox Log-rank analysis was used to test for differences in survival rates across all treatment groups and when significant, survival between a single clothianidin concentration and the 0ppb control.

### Test procedure for RNAseq analysis

To test the haploid susceptibility hypothesis that males have fewer gene products available to detoxify clothianidin than workers, RNAseq analysis was used to conduct genome-wide transcriptome analysis of workers and males after consumption 75uL of either the 5ppb test solution (0.374ng/bee; 3 workers and 3 males) or 1.4% DMSO in stock sugar solution (vehicle control; 3 workers and 3 males) once a day for 5 consecutive days. This clothianidin dose was selected because it did not reduce worker and male survival in our initial series of feeding experiments.

#### Treatment and sample preparation

Individuals were euthanized at -20°C 24 hours after the last dose (day 6), immediately dissected on ice by removing legs, wings, and proboscis, and transferred to -80°C until ready for extraction. (This entire process was performed as quickly as possible to minimize RNA degradation, usually completed within 5 minutes.) Individual sample extractions were completed in an RNA-specified biosafety cabinet by crushing the bee body through pestle homogenization in Trizol (ThermoFischer Scientific, USA) following the suppliers recommendations for RNA isolation followed by chloroform phase separation. The aqueous phase containing RNA was collected, then purified using a PureLink RNA Mini Kit (ThermoFischer Scientific, USA), ending with a 30 uL elution step. Concentration and purity (A260/280 and A260/230) were measured using Nanodrop. Purified RNA was stored at -80°C until use.

#### RNA Sequencing

Purified RNA samples were diluted to 100ng/uL in sterile, DNase-free, RNase-free water and then shipped to Quick Biology (Pasadena, USA) for processing. Libraries for RNA-Seq were prepared with a KAPA Stranded RNA-Seq Kit. The workflow consisted of mRNA enrichment, cDNA generation, and end repair to generate blunt ends, A-tailing, adaptor ligation and PCR amplification. Different adaptors were used for multiplexing samples in one lane. Sequencing was performed on Illumina Hiseq3000/4000 for a pair end 150 run. Data quality check was done on an Illumina SAV (San Diego, USA). De-multiplexing was performed with Illumina Bcl2fastq2 v 2.17 program. Reads were first mapped to the latest UCSC transcript set using STAR version 2.4.1d and the gene expression level was quantified using an E/M algorithm in Partek Genomics Suite (Partek, Inc. USA). Gene expression levels were normalized to total counts.

All four experimental groups produced a similar number of mean total reads: worker controls 4.951*10^6^ ± 1.845*10^5^, worker clothianidin 5.147*10^6^ ± 4.004*10^5^, male controls 5.520*10^6^ ± 8.654*10^4^, and male clothianidin 5.889*10^6^ ± 1.168*10^5^. Reads were aligned to the BIMP_2.0 genome [43], resulting a high mean percentage of alignment in all four groups: worker control 73.20% ± 3.35, worker clothianidin 80.41% ± 1.573, male control 79.15% ± 1.511, male clothianidin 77.58% ± 1.938, corresponding with 15,896 identified genes. RNA sequencing data quality and alignment for sex and treatment are shown in Table S4.

#### Statistical Analysis

Normalized gene expression levels were compared between groups using the differential gene expression (GSA) algorithm in Partek Genomics Suite (Partek, Inc., USA). To initially isolate all putative detoxification genes induced by clothianidin consumption, test and vehicle control gene expression patterns were compared separately for each sex. A gene was considered to be differentially expressed if comparisons yielded both a p-value less than 0.05 and a fold change greater than 1.5. All differentially expressed genes in workers and males were then pooled to create a list of genes with clothianidin-induced expression.

To test for sex differences in the expression level of pooled clothianidin-induced genes, expression levels were then compared between workers and males in the test group (bees chronically exposed to clothianidin at concentration of 5ppb). For this comparison, a gene was considered to have sex-specific expression if P<0.05. Genes with this significance threshold were then categorized as being involved in either Phase I (oxidation, hydrolysis and reduction) or Phase II (conjugation with sugars, glutathione, amino acids, etc.) detoxification or assigned to some other biological process. This method of identifying and categorizing putative detoxification genes has been used previously in bumblebees and other insects [25] and [44]. Genes were assigned a particular biological processes based on information from the UniProt online gene database (http://www.uniprot.org/).

## General

We thank C. Emery and J. Letourneau for assistance with feeding and data collection.

## Supporting Information

**Fig S1. Schematic of setup for individual feeding assay.** Bees were isolated from the colony and housed in a 16oz plastic container with a feeding cup positioned directly under an open vent in the screen lid. Test sugar solutions were delivered through a hole in the lid with a micropipette. Individuals were supplied with pollen *ad libitum*.

**Table S1. Percent of alive queens, workers and males consuming test solutions (clothianidin concentration given in ppb) over the 7-day testing period.**

**Table S2. List of clothianidin-induced non-detoxification genes with increased expression in worker (W) and male (M) *Bombus impatiens*.** Biological process known to be associated with each gene is also shown.

**Table S3. List of clothianidin-induced genes showing a similar expression level in worker (W) and male (M) *Bombus impatiens***.

**Table S4. RNA sequencing results and quality control**.

